# Composition fluctuations in lipid bilayers

**DOI:** 10.1101/078030

**Authors:** Svetlana Baoukina, Dmitri Rozmanov, D. Peter Tieleman

**Affiliations:** Centre for Molecular Simulation and Department of Biological Sciences, 2500 University Dr. NW, Calgary AB T2N 1N4, Canada; Department of Information Technologies, Research Computing Services, University of Calgary 2500 University Dr. NW, Calgary AB T2N 1N4, Canada

## Abstract

Lipid bilayers constitute the basis of biological membranes. Understanding lipid mixing and phase behavior can provide important insights into membrane lateral organization (the “raft” hypothesis). Here we investigate model lipid bilayers below and above their miscibility transition temperatures. Molecular dynamics simulations with the MARTINI coarse-grained force field are employed to model bilayers on a length scale approaching 100 nm and a time scale of tens of microseconds. Using a binary mixture of saturated and unsaturated lipids, and a ternary mixture of a saturated lipid, an unsaturated lipid and cholesterol we reproduce the coexistence of liquid-crystalline and gel, as well as liquid-ordered and liquid-disordered phases. By raising the temperature or adding hybrid lipids (with a saturated and an unsaturated chain), we induce a gradual transition from a two-phase to a one-phase state. We characterize the evolution of bilayer properties along this transition. Domains of coexisting phases change to dynamic heterogeneity with local ordering and compositional de-mixing. We analyze the structural and dynamic properties of domains, sizes and lifetimes of composition fluctuations, and calculate the in-plane structure factors.

## Introduction

Lipid bilayers constitute the basis for cell membranes. They contain multiple lipid species, which are distributed non-uniformly along the bilayer plane (1). This distribution modulates the bilayer properties and creates an optimized environment for protein function. The "raft" hypothesis suggests that membrane components exhibit nano-scale dynamic heterogeneity (2). Rafts are enriched in saturated lipids and cholesterol and incorporate selected proteins (3). They have sizes in the range of 10-200 nm and short lifetimes (10^−3^-10^0^ s). Rafts are believed to be important in membrane trafficking, signal transduction, and entry of pathogens (4–7). However, direct experimental observation of rafts is challenging due to their dynamic nano-scale nature. Rafts are believed to represent nano-domains of the liquid-ordered (Lo) phase in the liquid-disordered (Ld) phase. Yet the small size and short lifetime of these Lo domains cannot be explained by classical theories for phase separation. As a result, the existence of rafts and mechanisms of their formation remain elusive (8–10).

The difficulty in characterizing the lateral organization of cell membranes lies in their complexity. A crowded environment with many lipid and protein players, it is coupled to the cytoskeleton and affected by active cellular processes (11). Characterizing phase behavior of lipids alone requires building multi-dimensional phase diagrams (12). Numerous membrane proteins interact preferentially with different lipids, and vary in size, membrane partitioning and mobility. Lipid transport interferes kinetically with these interactions, and coupling to cytoskeleton bounds them spatially. Many theories describing these phenomena attempt to explain raft formation (see e.g. (11, 13, 14) for review). Yet distinguishing between these different theories is difficult, in part due to a lack of understanding of lipid phase behavior.

Lipid-lipid interactions alone could produce rafts via several different mechanisms. The first group of mechanisms is based on phase separation with limited growth of domains. Domain growth could be limited due to inter-domain repulsion. Repulsion may be caused by uncompensated headgroup dipoles in the domains (15) or from domain curvature (16). A large number of small domains could become favorable (due to entropy gain) at low line tension at phase boundaries (17). Line tension may be lowered by linactants such as hybrid lipids. The second group of mechanisms is based on dynamic heterogeneity with local structure and order (18). Dynamic heterogeneity develops in one phase due to lateral density and composition fluctuations. Fluctuations increase in magnitude upon approaching a phase transition and become extremely strong in the vicinity of a critical point (19). Rafts could be a manifestation of two-dimensional micro-emulsion – a liquid with local structure and a tendency to order (20).

Substantial progress in understanding lipid phase behavior has been made by studying lipid bilayers of simple composition. The coexistence of macroscopic domains of the Lo and Ld phases has been experimentally reproduced in mixtures of saturated and unsaturated lipids and cholesterol (see (21, 22) for review). Advanced methods provided details on the properties of coexisting Lo and Ld phases (23, 24). Transition from macro- to nano-scale Lo domains has been observed upon substitution of unsaturated lipids by so-called hybrid lipids with one saturated and one unsaturated chain (25, 26). In these studies, nano-domains are detected indirectly, and it is difficult to establish their origin due to limits in achievable imaging resolution. Distinguishing between nano-scale domains and fluctuations in experiments remains a challenging task.

Here, we characterize the properties of composition fluctuations and domains of coexisting phases at the molecular level. To this end, we simulate model lipid bilayers above and below their miscibility transition temperature. We employ molecular dynamics (MD) simulations with the Martini coarse-grained force field (27). The Martini force field has been used in many studies of phase separation in model lipid bilayers (see e.g. (28, 29) for review). It has recently been used to study the properties of complex lipid bilayers in a liquid phase mimicking the composition of real plasma membranes (30). Here we focus on the transition from a one-phase to a two-phase state in simple lipid mixtures. We consider two types of phase coexistence relevant for cell membranes: liquid-liquid and liquid-solid, and change the bilayer phase behavior by varying the temperature and adding a hybrid lipid.

## Methods

We performed Molecular dynamics (MD) simulations with the Gromacs software package (v.4.6.5) (31). Lipid bilayers were simulated with the Martini coarse-grained (CG) force field (27). We used a binary mixture of dipalmitoyl-phosphatidylcholine (DPPC) and dilinoleoyl-phosphatidylcholine (DUPC) to model the coexistence of the liquid-crystalline (L*α*) and gel (L*β*) phases, and a ternary mixture of DPPC, DUPC and cholesterol to model the coexistence of the liquid-disordered (Ld) and liquid-ordered (Lo) phases. The molar ratios for the two mixtures were 3:2, and 7:7:6, respectively. These mixtures were simulated in a temperature range of 270-340 K. At higher temperatures, the bilayers mixed and formed a single phase, in which the composition fluctuations were studied. At lower temperatures, the bilayers de-mixed and separated into domains of coexisting phases. A so-called hybrid lipid with one saturated and one unsaturated chain, palmitoyl-linoleoyl-phosphatidylcholine (PUPC), was added to the two mixtures in the following way: a specific fraction of the DUPC lipids in each leaflet was randomly replaced by the PUPC lipid, and the bilayer was equilibrated at a T=340 K. Thus obtained molar ratios were DUPC: PUPC 8:2, 7:3 and 6:4 in the L*α*-gel mixture (20, 30 and 40 % PUPC substituting DUPC), and 6:1, 5:2, and 4:3 in the Ld-Lo mixture (14, 29 and 43 % PUPC substituting DUPC). These concentrations of the hybrid lipid were selected to achieve noticeable effects on phase separation and composition fluctuations, and, at the same time, to maintain phase coexistence in the selected interval of temperatures.

In the Martini force field, molecules are represented by particles that group approximately four heavy atoms together. All the lipids are standard components of the force field. The system setup consisted of a randomly mixed bilayer in water. Two system sizes were used: smaller bilayers (~35×35×15 nm^3^) contained 4608 lipids and were solvated in 128,000 CG water particles; larger bilayers (~70×70×30 nm^3^) contained 18432 lipids and were solvated in 1,024,000 CG water particles. At lower temperatures (< 290 K), the anti-freeze water particles substituted ~ 5% of regular water particles to prevent water crystallization, which is a standard practice in the Martini force field (27). For non-bonded interactions, the standard cutoffs for the MARTINI force field were used: the Lennard-Jones potential was shifted to zero between 0.9 and 1.2 nm; the Coulomb potential was shifted to zero between 0 and 1.2 nm with a relative dielectric constant of 15. The time step was 20 fs with neighbor list updates every 10 steps. Lipids and water were coupled separately to a target temperature using the velocity rescaling thermostat (32) with a time constant of 1 ps. The normal and tangential pressures of 1 bar were maintained using the Berendsen barostat(33) with the semi-isotropic coupling scheme, a time constant of 4 ps and compressibility 5⋅10^−5^bar^−1^. The simulation time was 10-30 *μ*s; longer times corresponded to the cases near or with phase separation. The last microsecond of the trajectory was used for analysis.

Quantitative analysis of composition fluctuations and domains of coexisting phases was performed using a combination of custom scripts. The areas per lipid were defined based on Voronoi tessellation using a Matlab program (v. R2014b). Lateral heterogeneity was analyzed based on local lipid environment (34), defined as the first surrounding shell of nearest neighbors. The Voronoi sites with a local concentration of DPPC and cholesterol above the average concentration in the bilayer were assigned to an "ordered cluster". The ordered clusters were then grouped together using the connectivity matrix. In these grouped clusters, the distinction between composition fluctuations and domains of coexisting phases was made based on structural and dynamic properties. The boundary was calculated as the sum of Voronoi edges between the cluster and its surroundings. The overlap (registration) of the clusters between the leaflets was calculated as the area of the clusters aligned (i.e. in register) in the two leaflets divided by to the total area of the clusters in the leaflet.

The in-plane two-dimensional (2D) radial distribution function (RDF) for the bilayers was calculated as the average ratio of the lipid density at the distance r from the center of mass of the lipid molecule to the average density in the bilayer. In these calculations, we considered the unsaturated lipid only, as it is enriched in the disordered phase in all cases, without long-range translational order (which avoids additional undulations on RDFs corresponding to intermolecular distances). The correlation length, *ξ*, was calculated from an exponential fit to the 2D RDFs using the formula:

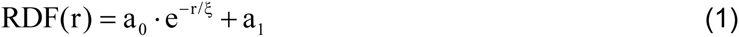

The correlation time, *τ*, was calculated from the time decay of the local density time correlation function using a single exponential fit as in (1). The local lipid density was sampled on a 2D grid of 20×20 cells from 10 consecutive trajectory time frames with the time step of 10 ps. It was assumed that such time is short relative to time scale of diffusion in the system so that the changes in the density are acceptably small.

The chain orientational order parameter, S_z_, was calculated using the formula:

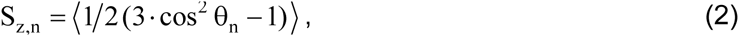

where *θ*_n_ is the angle between the vector connecting the n−1 and n+1 sites of the hydrocarbon chain and the monolayer normal z averaged over all sites for both chains and over all lipids, except for cholesterol. The mean curvature of the bilayer:

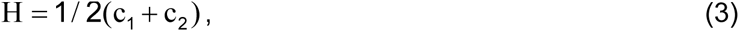

was obtained from the principal curvatures, c_1_ and c_2_ as in previous studies (35). The in-plane 2D static structure factor, 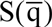, was calculated from the scattering of the molecular centers of mass using the formula:

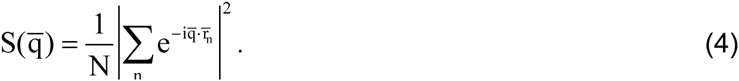

Here the scattering length of all scattering centers, N, was assumed constant, 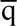 is the wave vector, and 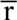 is the real space vector. The calculated structure factor as a function of 2D wave vector was then averaged over the polar angle to give the radial component, S(q).

## Results

### Temperature dependence of the bilayer properties

We simulated lipid bilayers composed of mixtures of DPPC: DUPC in ratio 3:2 and of DPPC: DUPC: cholesterol in ratio 7:7:6. At lower temperatures, the bilayers separated into the coexisting L*α* and gel, and Ld and Lo phases, respectively. Upon raising the temperature, the bilayers mixed to form a single phase. Characteristic snapshots of the phase behavior in the two mixtures are shown in Figures 1a,b and 2a,b, and a summary of all simulations is given in Table 1a,b.

**Table 1.** Bilayer properties.

| T, K | $A_{L_2}$, nm<sup>2</sup> | $A_{DPPC}$, nm<sup>2</sup> | $A_{DUPC}$, nm<sup>2</sup> | $C_{DPPC}$ | $C_{DUPC}$ | $D_{DPPC}$, 10<sup>7</sup> cm<sup>2</sup>/s | $D_{DUPC}$, 10<sup>7</sup> cm<sup>2</sup>/s | $S_z$ | Phase |
| --- | --- | --- | --- | --- | --- | --- | --- | --- | --- |
| 340 | 0.682<br>±0.004 | 0.674<br>±0.005 | 0.740<br>±0.013 | 0.88<br>±0.01 | 0.12<br>±0.01 | 4.1<br>±0.2 | 4.6<br>±0.1 | 0.30<br>±0.02 | Lα |
| 320 | 0.652<br>±0.004 | 0.645<br>±0.004 | 0.712<br>±0.012 | 0.89<br>±0.01 | 0.11<br>±0.01 | 3.0<br>±0.1 | 3.1<br>±0.1 | 0.33<br>±0.03 | Lα |
| 300 | 0.621<br>±0.003 | 0.614<br>±0.004 | 0.685<br>±0.009 | 0.91<br>±0.01 | 0.09<br>±0.01 | 1.7<br>±0.1 | 1.9<br>±0.2 | 0.35<br>±0.02 | Lα |
| 290 | 0.603<br>±0.003 | 0.598<br>±0.003 | 0.665<br>±0.012 | 0.92<br>±0.01 | 0.08<br>±0.01 | 1.3<br>±0.1 | 1.4<br>±0.1 | 0.40<br>±0.04 | Lα |
| 285 | 0.593<br>±0.003 | 0.587<br>±0.003 | 0.661<br>±0.012 | 0.93<br>±0.01 | 0.07<br>±0.01 | 1.0<br>±0.1 | 1.1<br>±0.1 | 0.42<br>±0.04 | Lα |
| 280 | 0.468<br>±0.001 | 0.467<br>±0.001 | 0.565<br>±0.031 | 0.99<br>±0.01 | 0.01<br>±0.01 | 0.05<br>±0.01 | 0.7<br>±0.1 | 0.90<br>±0.01 | gel+Lα |
| 270 | 0.464<br>±0.001 | 0.463<br>±0.001 | 0.517<br>±0.019 | 0.99<br>±0.01 | 0.01<br>±0.01 | 0.02<br>±0.01 | 0.4<br>±0.1 | 0.91<br>±0.01 | gel+Lα |

**b.**
| T, K | $A_{L_2}$, nm<sup>2</sup> | $A_{DPPC}$, nm<sup>2</sup> | $A_{DUPC}$, nm<sup>2</sup> | $A_{CHOL}$, nm<sup>2</sup> | $C_{DPPC}$ | $C_{DUPC}$ | $C_{CHOL}$ | $D_{DPPC}$, 10<sup>7</sup> cm<sup>2</sup>/s | $D_{DUPC}$, 10<sup>7</sup> cm<sup>2</sup>/s | $S_z$ | Phase |
| --- | --- | --- | --- | --- | --- | --- | --- | --- | --- | --- | --- |
| 360 | 0.525<br>±0.006 | 0.623<br>±0.006 | 0.690<br>±0.013 | 0.310<br>±0.005 | 0.56<br>±0.01 | 0.10<br>±0.01 | 0.34<br>±0.01 | 3.3<br>±0.1 | 4.1<br>±0.2 | 0.33<br>±0.03 | Ld |
| 340 | 0.496<br>±0.007 | 0.593<br>±0.007 | 0.662<br>±0.012 | 0.296<br>±0.005 | 0.57<br>±0.01 | 0.08<br>±0.01 | 0.35<br>±0.01 | 2.0<br>±0.1 | 2.6<br>0.1 | 0.37<br>±0.02 | Ld |
| 320 | 0.459<br>±0.007 | 0.553<br>±0.006 | 0.625<br>±0.017 | 0.282<br>±0.005 | 0.57<br>±0.01 | 0.07<br>±0.01 | 0.36<br>±0.01 | 1.1<br>±0.2 | 1.5<br>±0.1 | 0.42<br>±0.03 | Ld |
| 310 | 0.436<br>±0.006 | 0.532<br>±0.005 | 0.612<br>±0.017 | 0.274<br>±0.003 | 0.58<br>±0.01 | 0.04<br>±0.01 | 0.38<br>±0.01 | 0.6<br>0.1 | 1.3<br>±0.1 | 0.51<br>±0.03 | Lo+Ld |
| 300 | 0.425<br>±0.005 | 0.519<br>±0.006 | 0.606<br>±0.025 | 0.269<br>±0.004 | 0.59<br>±0.01 | 0.02<br>±0.01 | 0.39<br>±0.01 | 0.4<br>±0.1 | 1.2<br>±0.1 | 0.60<br>±0.06 | Lo+Ld |
| 290 | 0.421<br>±0.004 | 0.516<br>±0.004 | 0.600<br>±0.019 | 0.270<br>±0.003 | 0.59<br>±0.01 | 0.02<br>±0.01 | 0.39<br>±0.01 | 0.15<br>±0.01 | 0.90<br>±0.02 | 0.67<br>±0.04 | Lo+Ld |
| 280 | 0.407<br>±0.002 | 0.500<br>±0.002 | 0.571<br>±0.018 | 0.262<br>±0.003 | 0.59<br>±0.01 | 0.02<br>±0.01 | 0.39<br>±0.01 | 0.08<br>±0.02 | 0.65<br>±0.01 | 0.68<br>±0.04 | Lo+Ld |
| 270 | 0.398<br>±0.003 | 0.490<br>±0.004 | 0.544<br>±0.021 | 0.257<br>±0.003 | 0.59<br>±0.01 | 0.01<br>±0.01 | 0.40<br>±0.01 | 0.02<br>±0.01 | 0.50<br>±0.07 | 0.72<br>±0.02 | Lo+Ld |

Data for the small DPPC:DUPC 3:2 bilayer forming L $\alpha$ -gel phases (a) and the small DPPC:DUPC:cholesterol 7:7:6 bilayer forming Ld-Lo phases (b). Here $A_L$ is the average area per lipid in the ordered clusters, $A_i$ are the areas per lipid components in the ordered clusters, and $C_i$ are the concentration of lipid components in the ordered clusters; $D_i$ are coefficients of lateral diffusion in the bilayers; $i$ =DPPC, DUPC (a,b), and cholesterol (b); $S_z$ is the orientational order parameter for Martini bonds, averaged over the hydrocarbon chains in the ordered clusters.

**Figure 1.**
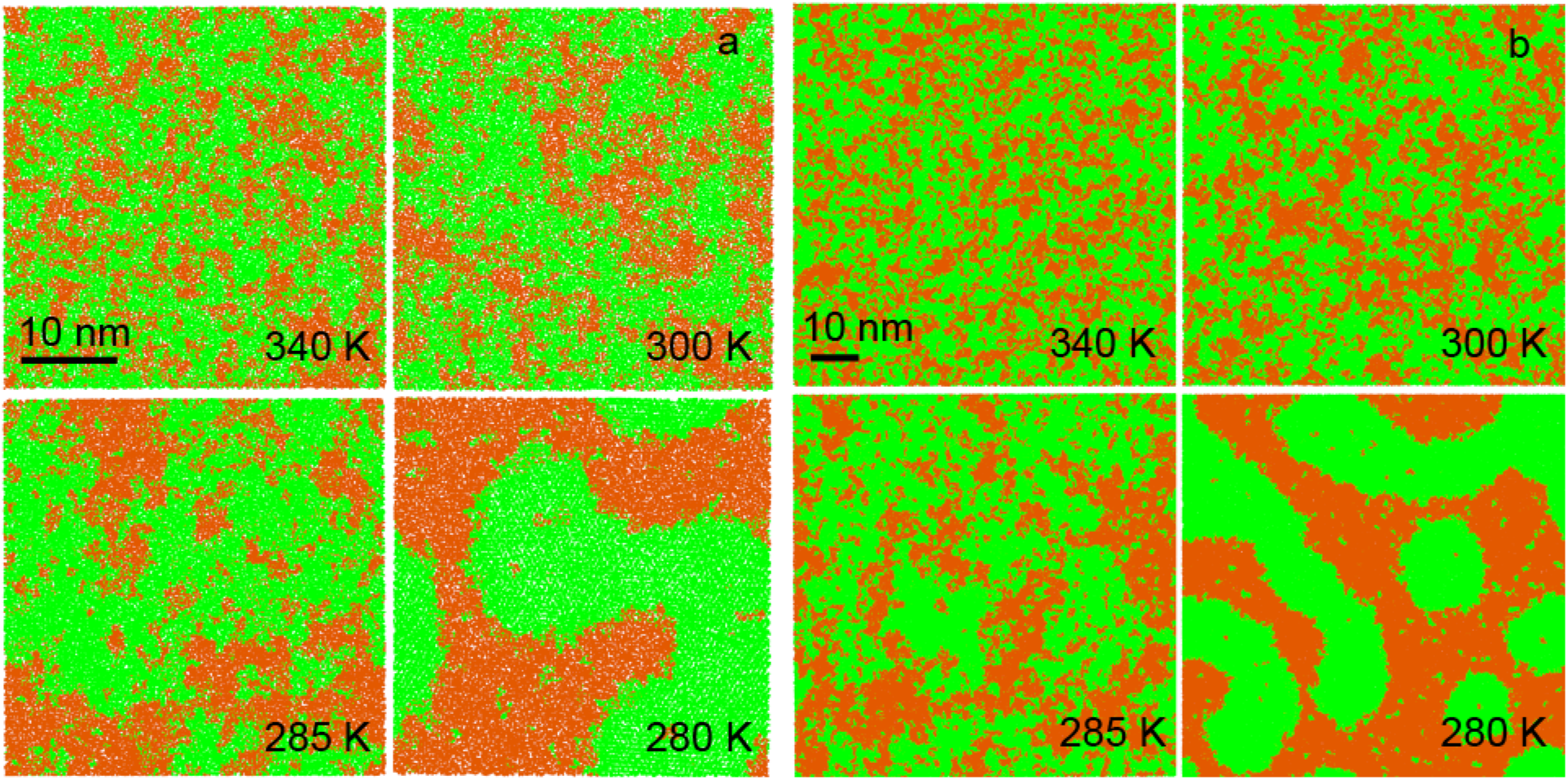
Phase behaviour of the DPPC: DUPC 3:2 small (a) and large (b) bilayers at selected temperatures. View from top, DPPC is shown in green, DUPC in orange, water not shown.

The first mixture of 3:2 DPPC: DUPC (Figure 1) forms the L*α* phase at 340 K. In this state, the bilayer is not homogeneous and contains small clusters of the two components. These clusters are manifestations of composition fluctuations. With decreasing temperature, the bilayer becomes more heterogeneous as the composition fluctuations grow. Domains of the gel phase appear in the surrounding L*α* phase at 280 K. A detailed view of the coexisting gel-L*α* phases is given in Figure 3a. The highly ordered gel phase contrasts with the disordered L*α* phase. Phase separation in this mixture occurs below the melting temperature of the saturated lipid (Tm~295 K in Martini (36)), which transforms into the gel phase. The segregation of the saturated and unsaturated lipids is driven by unfavorable interactions between the saturated and unsaturated lipid chains, which become more unfavorable as the saturated lipid transforms into the gel state (37). Transition to the gel phase is evident from an abrupt change of the average areas per lipid components, typical for a first order phase transition (Figure 4a). The transition is also characterized by significant changes in the lipid lateral diffusion coefficients and the chain orientational order (Table 1a). Near the de-mixing point (290 and 285 K) composition fluctuations become large in extent. The main phase transition in lipid bilayers is accompanied by strong fluctuations, being in a vicinity to a critical point (38, 39).

**Figure 2.**
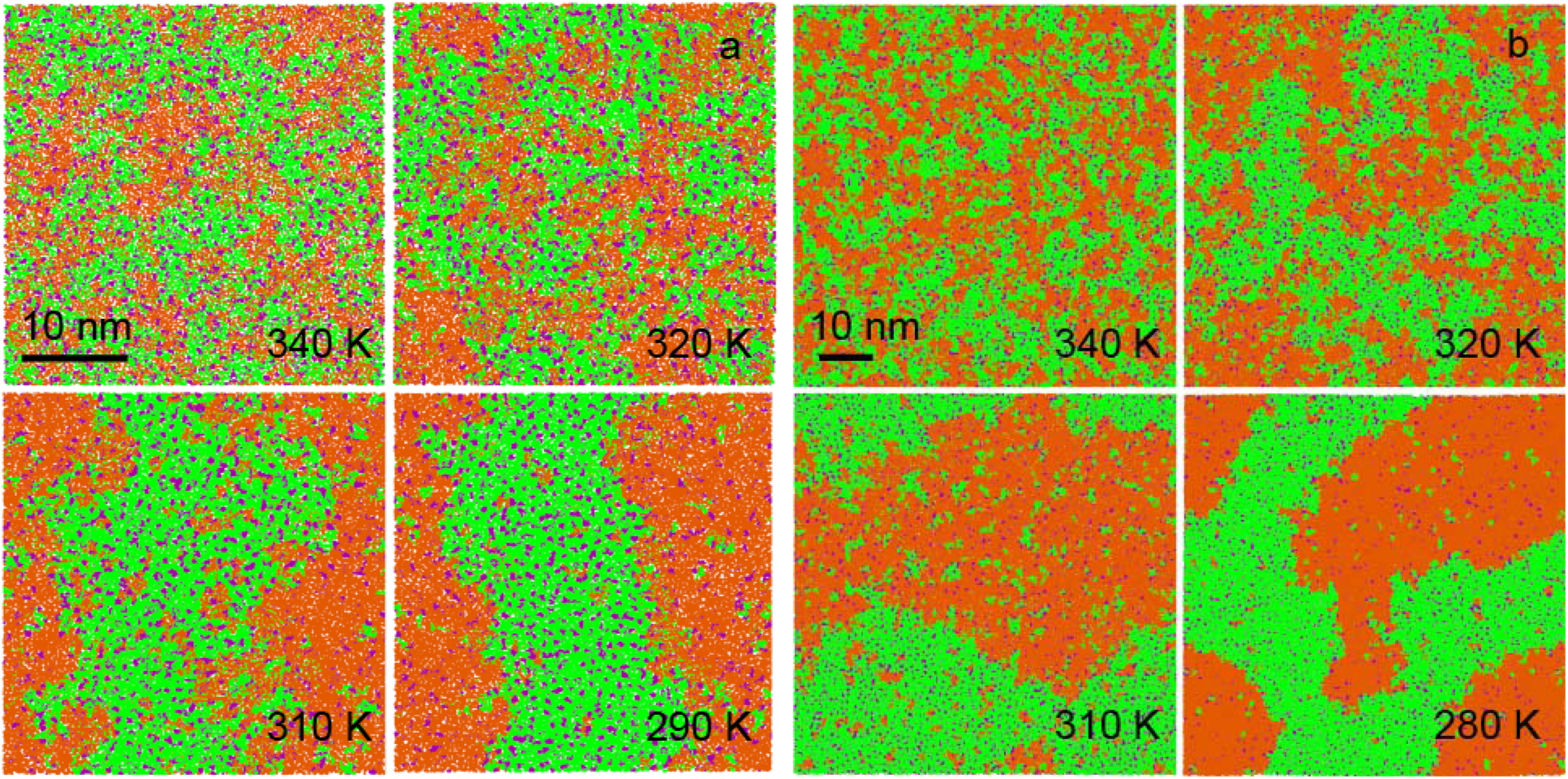
Phase behaviour of the DPPC: DUPC:cholesterol 7:7:6 small (a) and large (b) bilayers at selected temperatures. View from top, DPPC is shown in green, DUPC in orange, cholesterol in purple, water not shown.

**Figure 4.**
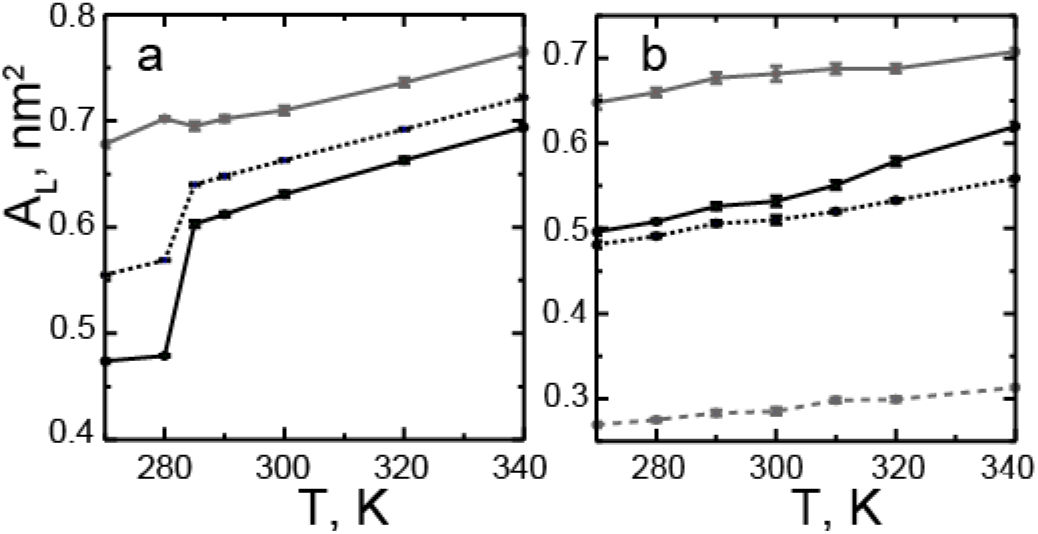
Areas per lipid averaged over the bilayer as a function of temperature for the DPPC: DUPC 3:2 (a) and DPPC: DUPC: cholesterol 7:7:6 (b) mixtures, data for small bilayers. DPPC area is shown as solid black, DUPC as solid grey, cholesterol as dashed gray, and the average area as dotted black.

The second mixture of 7:7:6 DPPC: DUPC: cholesterol (Figure 2) forms the Ld phase at 340 K. It contains small clusters (enriched either in DPPC and cholesterol or in DUPC), which increase in size at 320 K. At 310 K, the bilayer separates into Lo and Ld phases. Phase separation is induced by preferential interactions between the saturated lipid and cholesterol, leading to their segregation from the unsaturated lipid (37). A detailed view on the coexistence of the Lo and Ld phases is presented in Figure 3b. It shows that the phases noticeably differ in order, and the bilayer surface is bent at the phase boundary (see below). In contrast to the L*α*-gel mixture, strong composition fluctuations are absent, and the phase transition is continuous. The average areas per lipid, diffusion coefficients, and chain orientational order vary gradually as the temperature decreases (Figure 4b, Table 1b). In the absence of abrupt changes in these properties, the exact phase state of the bilayer can be determined from additional analysis.

**Figure 3.**
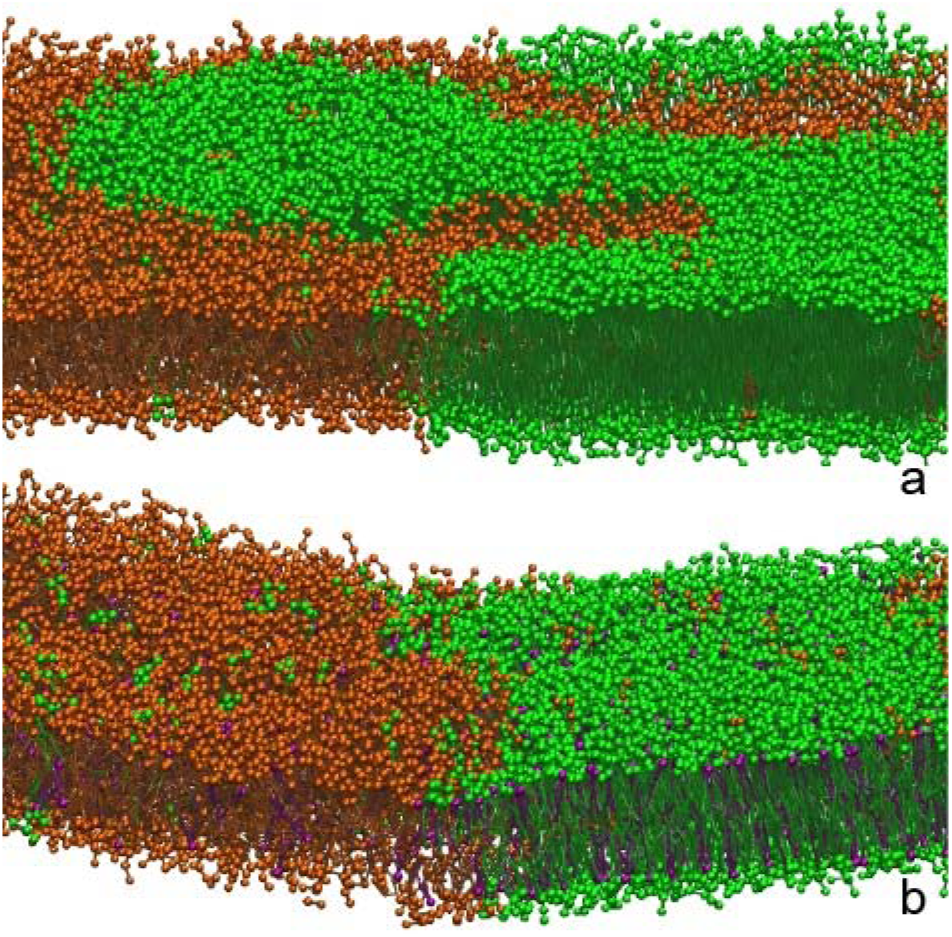
Detailed view on the coexistence of gel-L*α* (a) and Lo-Ld phases (b). The DPPC: DUPC 3:2 mixture at 280 K and the DPPC: DUPC:cholesterol 7:7:6 mixture at 290 K are shown.

Composition fluctuations can be distinguished from domains of coexisting phases by analyzing the in-plane (2D) radial distribution function (RDF) (Figure 4a,b). For one-phase bilayers, the RDFs decay exponentially as the correlations in density are short-range; for phase-separated bilayers, the RDFs decay is linear due to long-range density correlations in the domains (34, 40). Linear decay is followed by periodic undulations which correspond to the variation of densities of the lipid components between the two coexisting phases. As the temperature approaches the transition temperature, decay of RDFs occurs on a larger scale, in particular in the 3:2 DPPC: DUPC mixture at 285 and 290 K, where strong composition fluctuations are observed.

**Figure 5.**
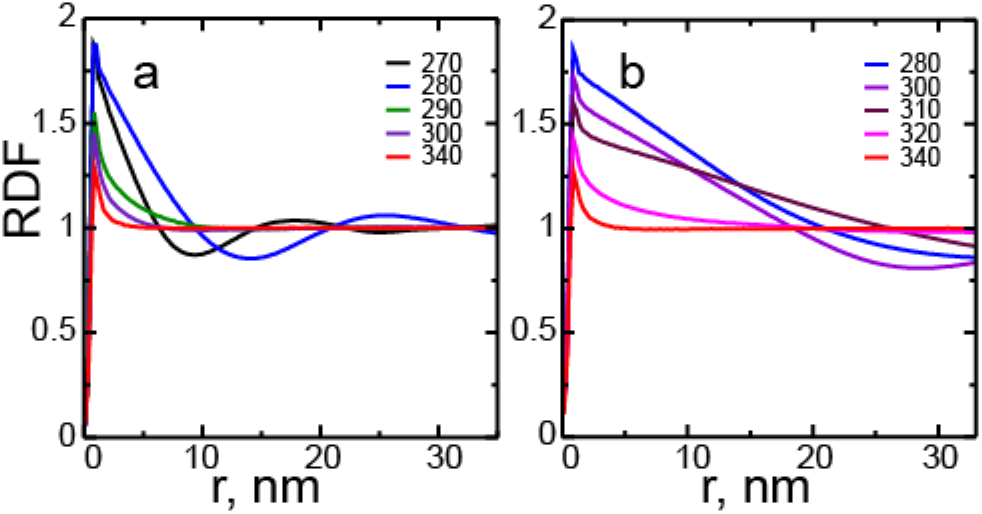
2D RDFs for the DPPC: DUPC 3:2 (a) and DPPC: DUPC: cholesterol 7:7:6 (b) large bilayers at different temperatures (K). RDFs are calculated for the centers of mass of DUPC lipids in the same leaflet.

**Figure 6.**
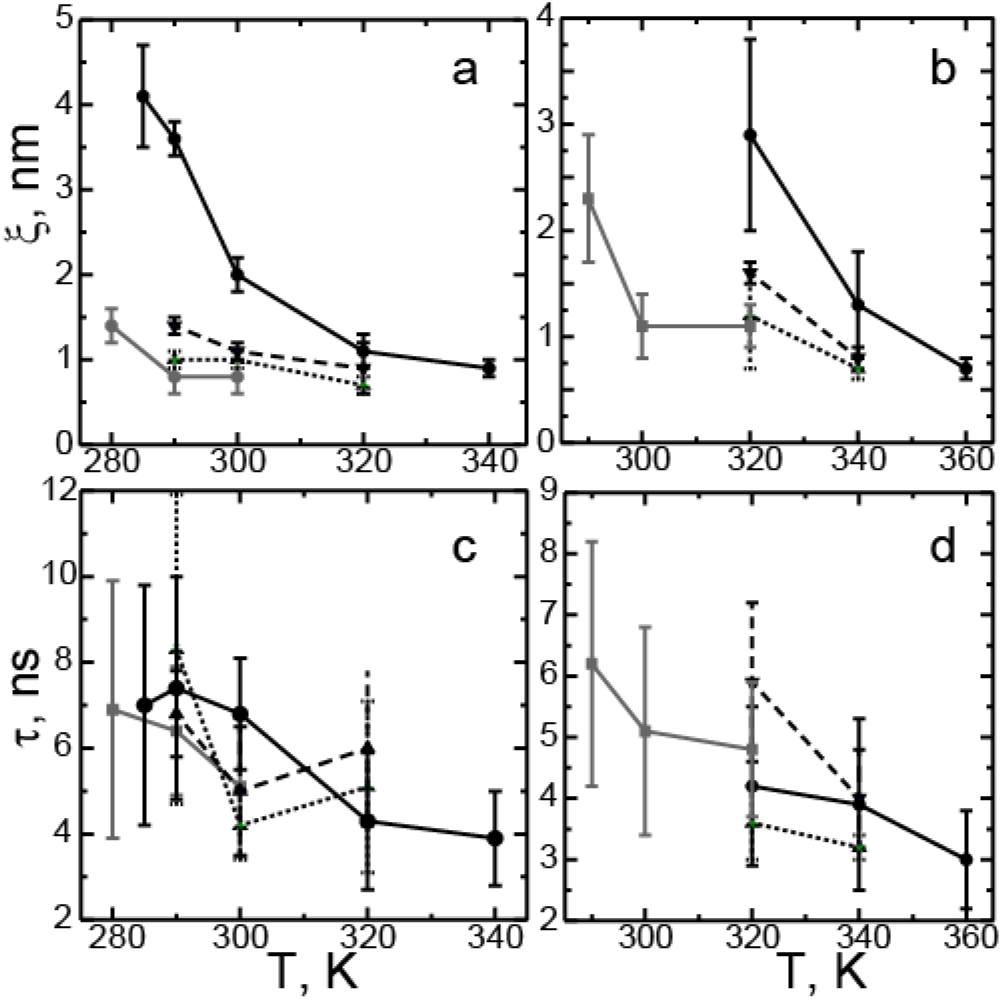
Correlation length(a,b) and time (c,d) for the DPPC: DUPC 3:2 (a,c) and DPPC: DUPC: cholesterol 7:7:6 (b,d) small bilayers. Different molar % of PUPC substituting DUPC are shown as follows: 0%, 20%, 30%, 40% in (a,c) or 0%, 14%, 29%, 43% in (b,d) as solid black, dashed black, dotted black, and solid grey, respectively.

To quantify composition fluctuations, we calculated the correlation lengths and times in one-phase bilayers (Figure 6a,b). As the temperature decreases and approaches the transition point, the correlation lengths increase as the fluctuations become stronger. The correlation times are expected to increase with increasing correlation length, but the data have a significant statistical uncertainty. The calculated correlation lengths and times are of the order of nm and ns, respectively. Nano-scale values are expected as the spatial extent of fluctuations is limited by the simulation box size (tens of nm).

**Figure 7.**
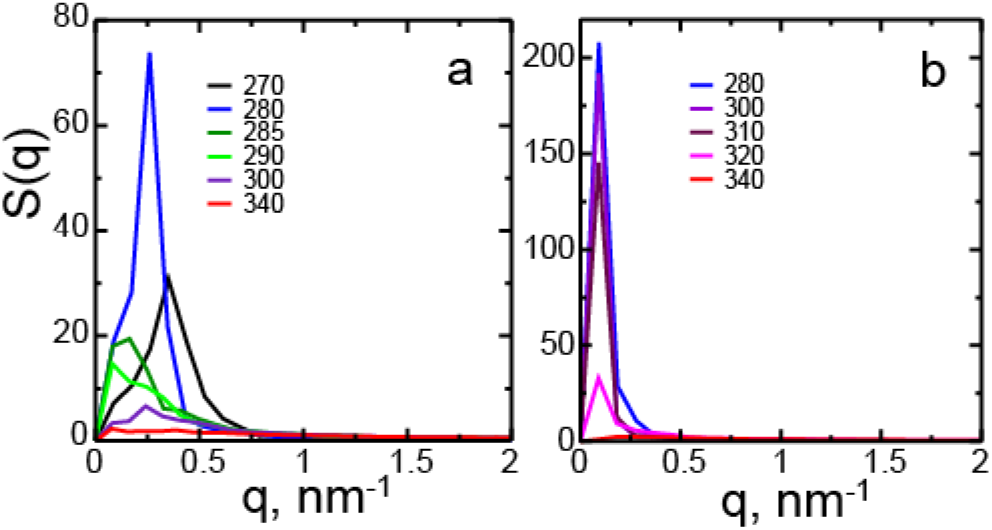
2D structure factors for the DPPC: DUPC 3:2 (a) and DPPC: DUPC: cholesterol 7:7:6 (b) large bilayers at different temperatures (K). Structure factors are calculated for the centers of mass of DUPC lipids in the same leaflet.

The lateral organization of the bilayers was also characterized by calculating the 2D structure factors (Figure 7a,b). Molecular centers of mass of the unsaturated lipid were used as the scattering centers. Note that the resolution of the structure factors in simulations is limited by the inverse simulation box size (~1/80 nm^−1^). The structure factors show large peaks in phase-separated bilayers. These peaks result from periodic variations of density in the coexisting phases, and correspond to a radial average of the domain spacing. In the L*α*-gel mixture, the peak is shifted towards larger wave vectors at 270 K (q~0.35 nm^−1^) compared to 280 K (q~0.26 nm^−1^) as the domains become smaller forming a thin network. Interestingly, strong composition fluctuations at 285 and 290 K manifest as intermediate peaks at similar wave vectors (q~0.16 nm^−1^). They result from strong correlations in space on large length scales comparable to domain size (tens of nm). Note that in the Ld-Lo mixture, the peaks of the structure factors are located at smaller wave vectors (q~0.09 nm^−1^), as the coexisting phases span the simulation box.

### Ordered clusters

To investigate the properties of composition fluctuations and domains of coexisting phases, we analyzed lipid clusters in each leaflet (see Methods). We clustered the sites with an increased local concentration of DPPC (and cholesterol in the ternary mixture), compared to their average concentration in the bilayer. Figure 8 shows that these clusters are more ordered as they have higher orientational order of lipid chains and smaller areas per lipid components compared to the bilayer averages (see also Table 1a,b). We thus call them the “ordered clusters” they represent domains of the gel or Lo phase in phase-separated bilayers, and composition fluctuations in one-phase bilayers. The area fraction of the ordered clusters increases with decreasing temperature; the more ordered phase domains have a larger area than fluctuations. In the 3:2 DPPC: DUPC mixture, the gel phase contains mainly DPPC. The properties of the gel phase are in agreement with previous simulation results (36) and experimental data (41). In the 7:7:6 DPPC: DUPC: cholesterol mixture, the concentration of DPPC and cholesterol in the Lo domains is nearly constant (~ 0.59 and 0.39, respectively), and is comparable to previously determined values for similar mixtures (42, 43). The average area per lipid in the Lo phase is in good agreement with experimental data (23). Based on diffusion coefficients, order parameters and areas per lipid, the Lo phase becomes substantially more ordered with decreasing temperature.

**Figure 8.**
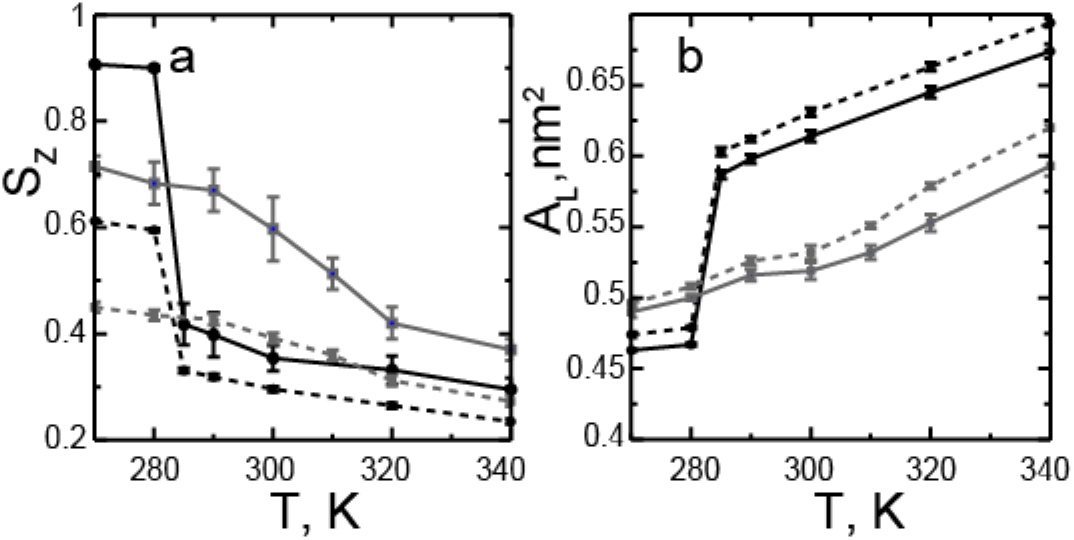
Chain orientational order parameter (a) and area per DPPC lipid (b) in the DPPC: DUPC 3:2 mixture (black) and the DPPC: DUPC: cholesterol 7:7:6 mixture (grey); solid lines correspond to the ordered clusters (see Methods), dashed lines to the averages in the bilayers.

**Figure 9.**
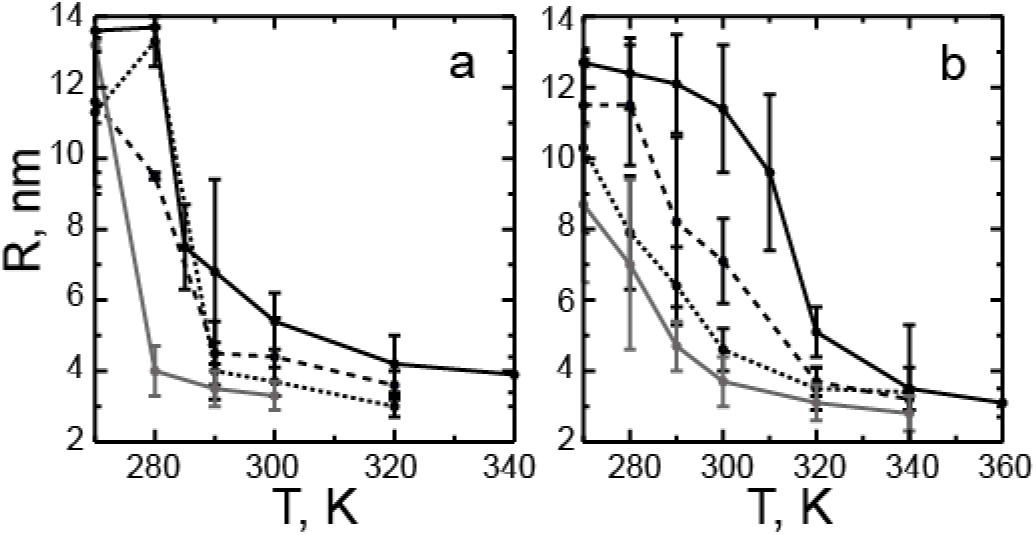
Radius of the ordered clusters as a function of temperature for the DPPC: DUPC 3:2 (a) and DPPC: DUPC: cholesterol 7:7:6 (b) small bilayers. Different molar % of PUPC substituting DUPC are shown as follows: 0%, 20%, 30%, 40% in (a) or 0%, 14%, 29%, 43% in (b) as solid black, dashed black, dotted black, and solid grey, respectively.

We then used the ordered clusters to analyze the differences between composition fluctuations and domains of coexisting phases. The bilayer phase state (i.e. one vs. two phases) was established based on the combination of 2D RDFs (linear vs. exponential decay, see above), areas per lipid, order parameters and diffusion coefficients. We found that fluctuations and domains noticeably differ in several ways. The cluster radius (Figure 9a,b) increases with decreasing temperature, and changes significantly on phase separation. Note that due to the limited simulation box size, the radii of both fluctuations and domains lie on the nano-scale, and it is not possible to estimate the final size of the domains (i.e. nano- vs. macro-). The boundary length (Figure 10a,b) is significantly larger for composition fluctuations compared to domains, in agreement with previous findings (34, 43). Here the boundary length was normalized by the perimeter of the circular cluster of the same area (a perfectly round cluster would thus have the boundary length of unity). Values much larger than unity indicate that fluctuations have a rough, irregular shape compared to domains.

**Figure 10.**
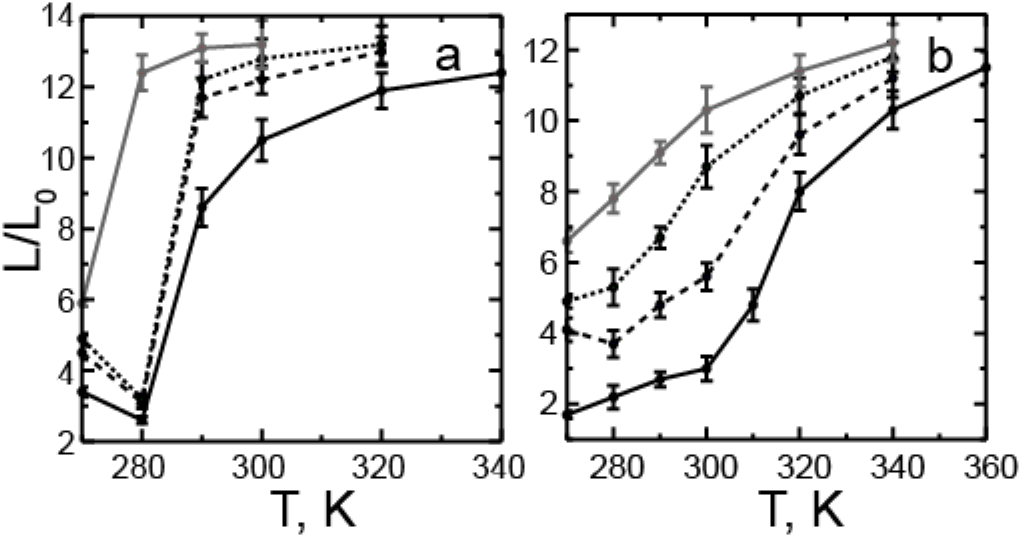
Boundary length of the ordered clusters as a function of temperature for the DPPC: DUPC 3:2 (a) and DPPC: DUPC: cholesterol 7:7:6 (b) small bilayers. The boundary length is normalized by the perimeter of a circular cluster of the same area. Different molar % of PUPC substituting DUPC are shown as follows: 0%, 20%, 30%, 40% in (a) or 0%, 14%, 29%, 43% in (b) as solid black, dashed black, dotted black, and solid grey, respectively.

**Figure 11.**
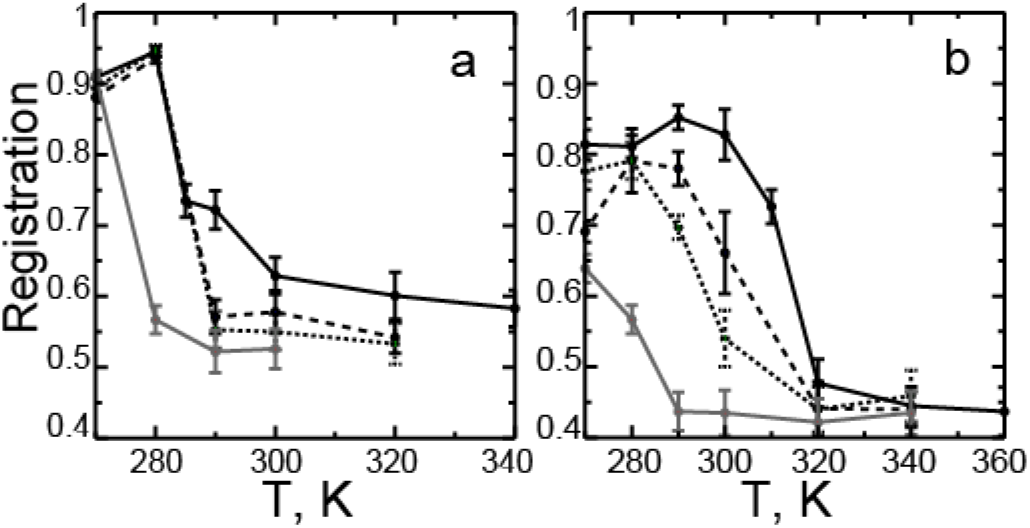
Overlap of the ordered clusters between the two leaflets as a function of temperature for the DPPC: DUPC 3:2 (a) and DPPC: DUPC: cholesterol 7:7:6 (b) small bilayers. Different molar % of PUPC substituting DUPC are shown as follows: 0%, 20%, 30%, 40% in (a) or 0%, 14%, 29%, 43% in (b) as solid black, dashed black, dotted black, and solid grey, respectively.

**Figure 12.**
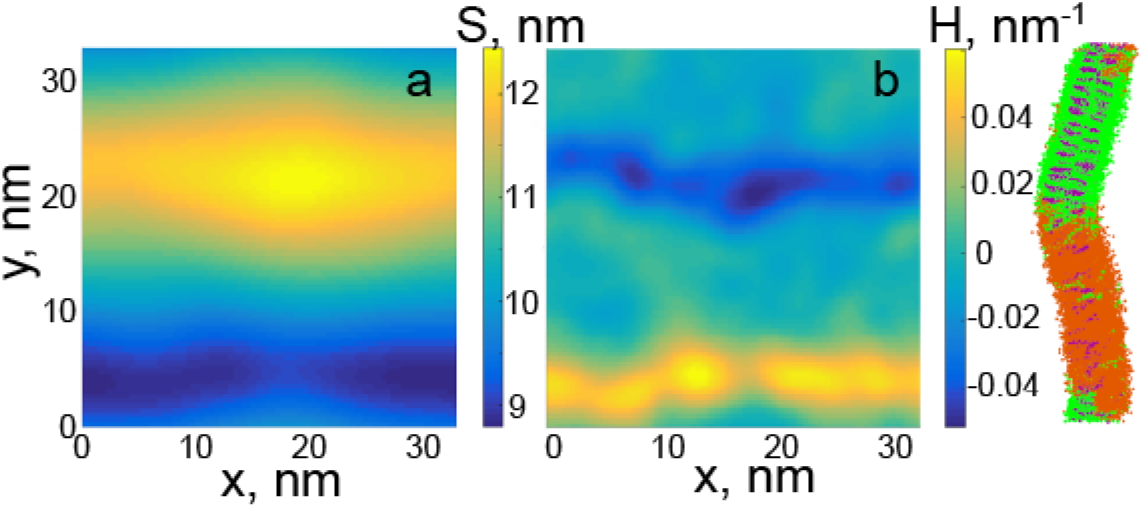
Curvature of DPPC: DUPC: cholesterol 7:7:6 bilayers. (a) Surface profile, S, and (b) mean curvature, H, are shown as functions of x and y coordinates in the bilayer plane. The right panel shows the corresponding side view of the bilayer; color scheme as in Figure 2.

The overlap (registration) between the two leaflets also differs significantly for domains and composition fluctuations (Figure 11). The overlap was quantified as the area fraction of the clusters aligned between the two leaflets. Complete overlap and complete anti-registration are expected to have values of 1 and 0, respectively. The overlap of uncorrelated clusters is expected to be between 0.57 and 0.50, as the area fraction of the ordered clusters in each leaflet lies between 0.31 (at higher temperatures) and 0.46 (at lower temperatures) (44). Therefore, inter-leaflet correlation of composition fluctuations can be considered negligible, except for strong fluctuations in the 3:2 DPPC: DUPC mixture at 285 and 290 K (overlap >0.7). In contrast, domains of coexisting phases overlap substantially. These difference in overlap between domains and fluctuations is consistent with the inter-leaflet 2D RDFs (see Figure S1 in Supporting Information) and in agreement with previous simulations (34, 43). Earlier simulations also showed that domain overlap depends on the thickness mismatch between the Lo and Ld phases, and increases with domain size (45). Macroscopic domains in symmetric bilayers are generally found in register in experimental studies (46–48).

In our simulations, overlap of Lo domains is smaller than of gel domains. Recent theoretical model predicts that incomplete registration of domains minimizes the deformation energy at the domain boundary, which reduces the line tension (49). This model assumes zero spontaneous curvatures of the leaflets. In our simulations, bilayers with the Ld-Lo phase coexistence are non-flat (see Figure 12) and have alternating regions of negative and positive curvature (with mean curvature ~ 0.05 nm^−1^). Curved symmetric bilayers with coexisting Ld and Lo phases were previously observed in simulations with the Martini model (45). The Lo phase in monolayers of similar composition in Martini model has a negative curvature (~ −0.06 nm^−1^) (35, 50). This is in agreement with experimental studies that suggest that the Lo phase has a negative spontaneous curvature, mainly resulting from a strongly negative curvature of cholesterol (24, 51). As the direction of bending is the opposite for the two leaflets, spontaneous curvature could not develop if the Lo domains overlapped completely. We hypothesize that incomplete overlap of Lo domains allows bending of symmetric bilayers as a result of spontaneous curvature of the leaflets.

### Effect of the hybrid lipid

We then substituted a fraction of the unsaturated lipid DUPC by the hybrid lipid PUPC in both mixtures (see Methods for details). Characteristic snapshots of the phase behavior for selected compositions are shown in Figures 13 and 14. A summary of all simulations is given in the Supporting Information (SI) in Tables S1 and S2.

The hybrid lipid generally induces mixing in both the L*α*-gel and the Ld-Lo bilayers, i.e. the phase transition temperature decreases (see Tables 1,2 and S1, S2). This behavior agrees with theoretical predictions (17, 52) and experimental observations (25). The distribution of PUPC is correlated with DUPC and inversely correlated with DPPC (see 2D RDFs in Figures S2, S3 in SI). These correlations are weak and the distribution of the hybrid lipid is nearly uniform in both mixtures in one-phase state. Approximately half of PUPC partitions into the ordered clusters, but only a small fraction of PUPC is present in the gel domains (see Tables S1, S2).

**Figure 13.**
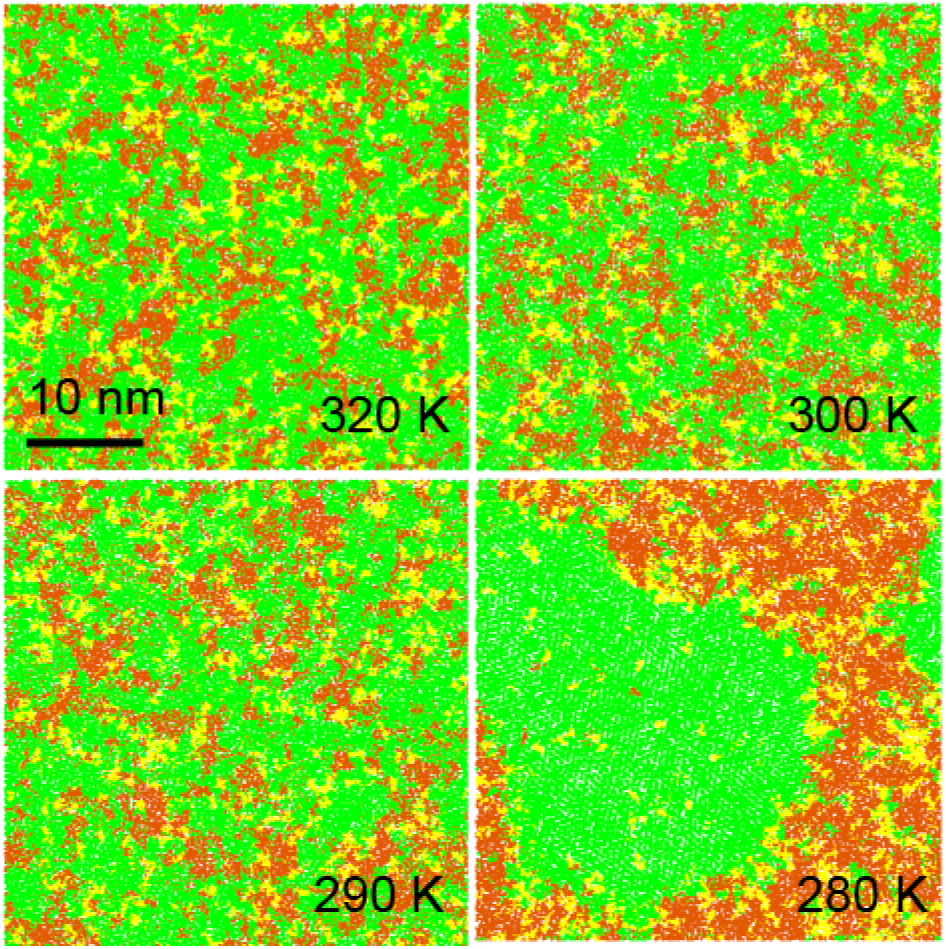
Phase behaviour of the DPPC: DUPC 3:2 small bilayers at selected temperatures, with 30% of DUPC substituted by PUPC. View from top, DPPC is shown in green, DUPC in orange, PUPC in yellow, water not shown.

**Figure 14.**
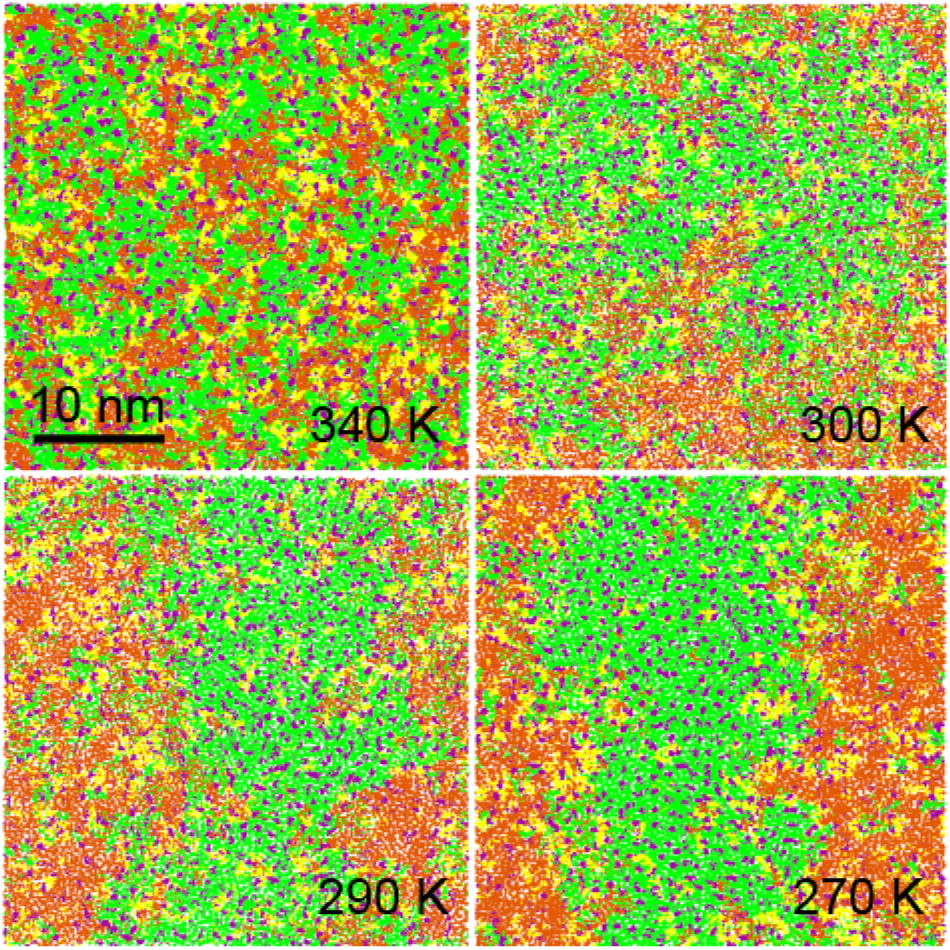
Phase behaviour of the DPPC: DUPC: cholesterol 7:7:6 small bilayers at selected temperatures, where 29% DUPC is substituted by PUPC. View from top, DPPC is shown in green, DUPC in orange, PUPC in yellow, cholesterol in purple, water not shown.

The properties of composition fluctuations and domains of coexisting phases are altered by the hybrid lipid. In the L*α*-gel mixture, strong composition fluctuations near the transition temperature are suppressed. Above the transition point (290 K at 20 and 30% of PUPC, and 280 K at 40% PUPC), the 2D RDFs decay on short scales (quantified by the correlation length, see below), and the peaks on the structure factor are small (compare Figures 7 and S4). In the Ld-Lo mixture, domains of the Lo phase appear more dynamic and disordered (Figure 14), and their boundary becomes more irregular (see below). The concentration of DPPC and, in particular cholesterol, in the Lo phase decreases.

With increasing concentration of the hybrid lipid, we observe the following trends in both mixtures. The correlation lengths of fluctuations (Figure 6a,b) decrease, in agreement with theoretical predictions (52). The correlation times (Figure 6c,d) approximately remain unchanged (given large statistical uncertainties). In other words, smaller fluctuations have longer lifetimes, which agrees qualitatively with recent theoretical predictions (53). Similar to the correlation length, the average radius of the ordered clusters decreases (see Figure 9), which is in qualitative agreement with experimental studies (where domain sizes decrease from micro- to nano-scale) (25, 26). In addition, the overlap of the clusters between the leaflets (Figure 11) becomes smaller, as was reported in earlier simulations (34).

**Figure 15.**
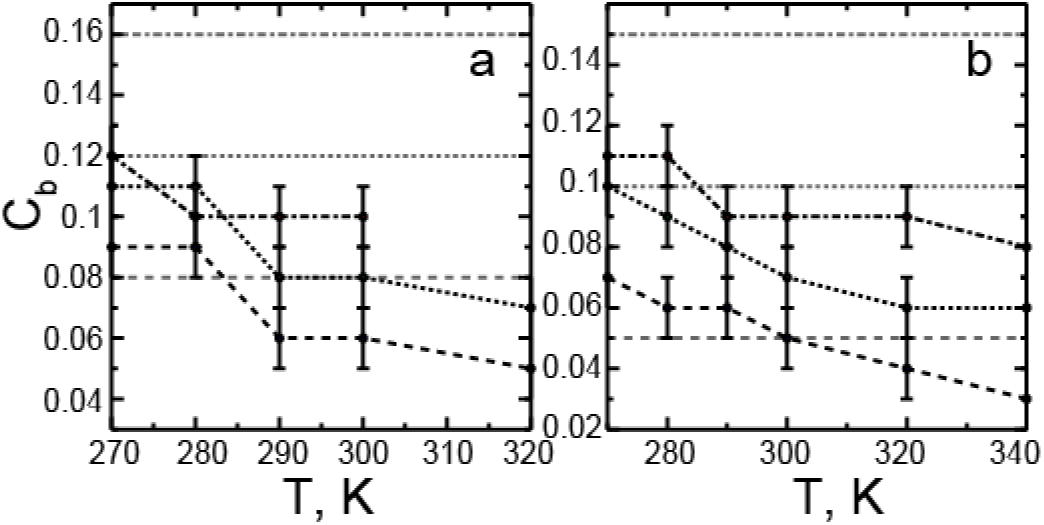
Concentration of the hybrid lipid PUPC (molar ratio) at the boundary as a function of temperature for the DPPC: DUPC 3:2 (a) and DPPC: DUPC: cholesterol 7:7:6 (b) small bilayers. Different molar % of PUPC substituting DUPC are shown as follows: 20%, 30%, 40% in (a) and 14%, 29%, 43% in (b) in black as dashed, dotted, and dashed-dotted lines, respectively; the average concentration of PUPC in the mixtures are shown in grey as dashed, dotted, and dashed-dotted lines for comparison.

The boundary length of domains and fluctuations (Figure 10) increases with increasing the concentration of the hybrid lipid. Theoretical model suggest that the hybrid lipid preferentially partitions at the phase boundary, reduces the line tension and thus favors domains of smaller size (17, 54). In this model the saturated chain of the hybrid lipid faces the saturated lipids enriched in the ordered phase, and the unsaturated chain faces the unsaturated lipids in the disordered phase. This alignment reduces the packing incompatibility and hydrophobic mismatch between the two phases, which lowers the free energy at the boundary. Previous simulations reported a small increase of the concentration of the hybrid lipid at the phase boundary (46, 55). In our simulations, the hybrid lipid does not show preferential partitioning to the boundary, its concentration at the boundary is comparable to or less than the average is the bilayer (Figure 15).

**Figure 16.**
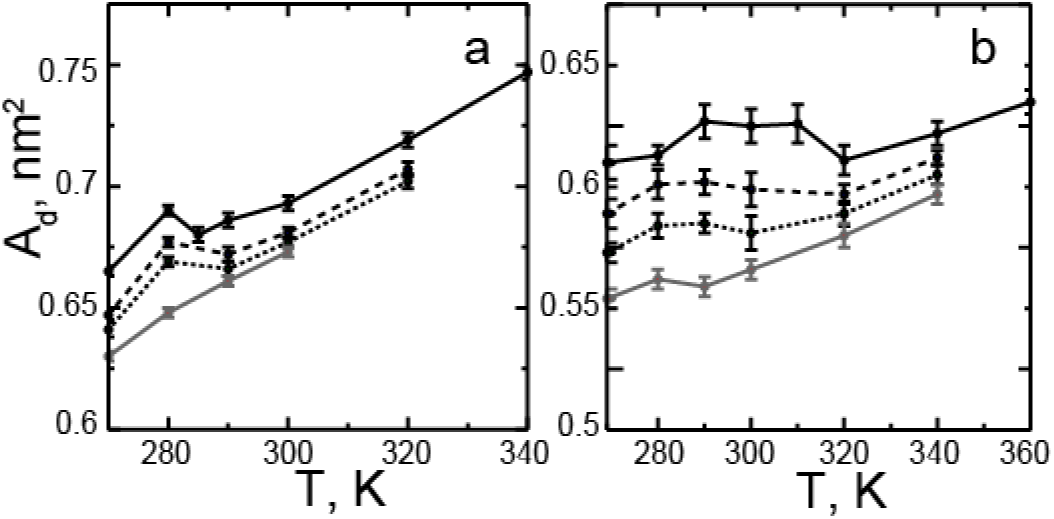
Average area per lipid in the disordered clusters for the DPPC: DUPC 3:2 (a) and DPPC: DUPC: cholesterol 7:7:6 (b) small bilayers. The clusters represent composition fluctuations or domains of the more disordered phase (L*α* or Ld). Different molar % of PUPC substituting DUPC are shown as follows: 0%, 20%, 30%, 40% in (a) and 0%, 14%, 29%, 43% in (b) as solid black, dashed black, dotted black, and solid grey, respectively.

Notably, the hybrid lipid changes the properties of the coexisting phases, in particular, the disordered phase. Figure 16 shows the average area per lipid in the disordered clusters, which correspond to either fluctuations enriched in the unsaturated lipid or domains of the disordered phase (L*α* or Ld). The area per lipid in the Ld phase changes non-monotonically with temperature. This property was found to decrease with increasing temperature in recent experimental studies (23), which was correlated with increasing concentration of cholesterol in the Ld phase. We observe only a small increase with temperature of cholesterol concentration in the Ld phase at a constant concentration of PUPC. Yet as the concentration of PUPC increases, the concentration of cholesterol in disordered clusters also increases. The area per lipid decreases substantially with increasing concentration of PUPC at all temperatures, in particular in the Ld-Lo mixture. The area changes are the opposite but less pronounced in the ordered clusters (see Tables S1, S2). The disordered clusters thus become more ordered and similar in properties to the ordered clusters. This decrease in the properties mismatch is expected to reduce the line tension at the boundary. Similar changes for the Ld/Lo phase coexistence with varying concentration of the hybrid lipid were observed in recent experimental studies (24).

## Discussion

We investigated the properties of composition fluctuations and domains of coexisting phases in lipid bilayers. The bilayers were simulated along the transition from a one-phase to a two-phase state. Two lipid mixtures with coexistence of either the L*α*/gel or the Ld/Lo phases were considered. The transition was induced by varying the temperature and lipid composition, where the unsaturated lipid was partially substituted by the hybrid lipid.

Above the transition temperature, lipid bilayers formed a heterogeneous liquid phase, and the degree of heterogeneity varied with temperature. At high temperatures, lipid mixtures were almost random containing only small dynamic clusters. As the temperature decreased approaching the phase transition temperature, characteristic size and lifetime of the clusters increased. These clusters are manifestations of density and composition fluctuations driven by more favorable interactions between specific lipid types, which vary with temperature (37). The fluctuations had the properties of the host phase (L*α* or Ld), i.e. were “homo-phase” in nature, in the sense that they were associated with the same phase state (56).

To quantify composition fluctuations, we analyzed the properties of the clusters with an increased local concentration of (DPPC, cholesterol) lipids forming a more ordered phase (gel or Lo). We found that the lateral density (inversed area per lipid) and chain orientational order in these clusters differed from their averages the bilayer, i.e. the clusters were more ordered. This local compositional de-mixing and ordering led to dynamic heterogeneity with local structure (18).

Composition fluctuations in one phase were distinct from domains of coexisting phases. They differed in the concentration of components, areas per lipid, order parameters, and diffusion coefficients. These properties changed abruptly in the 3:2 DPPC: DUPC mixture upon formation of the gel phase, but continuously in the 7:7:6 DPPC: DUPC: cholesterol mixture upon formation of the Lo phase. Generally, fluctuations were transient, while domains of coexisting phases were static, persistent in time and space.

The most significant differences between composition fluctuations and domains of coexisting phases were the following. First, composition fluctuations generally had negligible overlap between the two leaflets, while the overlap of domains of coexisting phases was substantial. Second, the average radius of fluctuations was much smaller compared to the radius of domains. Finally, the boundary length of composition fluctuations was substantially longer than that of domains of coexisting phases.

The hybrid lipid changed the properties of both composition fluctuations and domains of coexisting phases. The main changes were a reduction of the size of Lo domains and of fluctuations in both mixtures, and an increase of the boundary length. We found that these changes were related to ordering of the disordered clusters (domains), which thus became more similar in properties to the ordered clusters (domains). Preferential partitioning of the hybrid lipid to the boundary was not observed.

Here we considered simple lipid bilayers with fixed composition of several components. We believe that the findings of this work can be generalized to lipid bilayers with liquid-liquid and solid-liquid phase coexistence. We characterized the properties of nano-scale composition fluctuations and domains of coexisting phases. These structures lie on the length scale of rafts and are controlled only by lipid-lipid interactions. This study is thus important for understanding the role of lipids in the lateral organization of biological membranes.

## Conclusions

We simulated simple lipid bilayers with liquid-liquid and solid-liquid phase coexistence. A gradual transition from a two-phase to a one-phase state was induced by raising the temperature or adding a hybrid lipid. Along the transition, domains of coexisting phases change to dynamic heterogeneity with local ordering and compositional de-mixing. We identify significant differences between composition fluctuations and domains of coexisting phases, and characterize the effects of the hybrid lipid on fluctuations and domains.

## Acknowledgements

This work was supported by the Natural Sciences and Engineering Research Council (Canada). DPT is an Alberta Innovates Health Solutions Scientist and Alberta Innovates Technology Futures (AITF) Strategic Chair in (Bio)Molecular Simulation. Simulations were carried out on WestGrid/Compute Canada facilities.

## Supporting material

4 additional figures and 2 tables.

